# A classification-based approach to estimate the number of resting fMRI dynamic functional connectivity states

**DOI:** 10.1101/2020.06.24.161745

**Authors:** Debbrata K. Saha, Eswar Damaraju, Barnaly Rashid, Anees Abrol, Sergey M. Plis, Vince D. Calhoun

## Abstract

Recent work has focused on the study of dynamic (vs static) brain connectivity in resting fMRI data. In this work, we focus on temporal correlation between time courses extracted from coherent networks or components called functional network connectivity (FNC). Dynamic functional network connectivity (dFNC) is most commonly estimated using a sliding window-based approach to capture short periods of FNC change. These data are then clustered to estimate transient connectivity patterns or states. Determining the number of states is a challenging problem. The elbow criterion is a widely used approach to determine the optimal number of states. In our work, we present an alternative approach that evaluates classification (e.g. healthy controls versus patients) as a measure to select the optimal number of states (clusters). We apply different classification strategies to perform classification between healthy controls (HC) and patients with schizophrenia (SZ) for different numbers of states (i.e. varying the model order in the clustering algorithm). We compute cross-validated accuracy for different model orders to evaluate the classification performance. Our results are consistent with our earlier work which shows that overall accuracy improves when dynamic connectivity measures are used separately or in combination with static connectivity measures. Results also show that the optimal model order for classification is different from that using the standard k-means model selection method and that such optimization improves resulting in cross-validated accuracy. The optimal model order obtained from the proposed approach also gives significantly improved classification performance over the traditional model selection method. In sum, the observed results suggest that if one’s goal is to perform classification, using the proposed approach as a criterion for selecting the optimal number of states in dynamic connectivity analysis leads to improved accuracy in hold-out data.

## 1. Introduction

The unconstrained resting human brain has been shown to exhibit time-varying functional connectivity (FC) dynamics (Chang and Glover 2010, Hutchison et al. 2013, Sakoglu et al. 2010). Since then, researchers have developed several methods to estimate these intrinsic connectivity dynamics (Calhoun et al. 2014, Preti, Bolton, and Van De Ville 2017) and to study if these dynamic connectivity estimates provide additional diagnostic value above average (static) functional connectivity between brain regions that assumes constancy of connectivity throughout the scan (Damaraju et al. 2014, Rashid et al. 2016).

Functional connectivity among two brain regions in resting-state functional magnetic resonance imaging (fMRI) data is computed as a pairwise statistical dependency (usually correlation) between their time courses. To delineate brain functional connectivity, resting-state fMRI (rsfMRI) data has been analyzed using a variety of analytical tools. Two of the major approaches are (i) seed-based analysis (Biswal et al. 1995, Greicius et al. 2003) and (ii) data-driven methods, such as independent component analysis (ICA) (Calhoun et al. 2001b, Calhoun and Adali 2012, Calhoun, Eichele, and Pearlson 2009, Damoiseaux et al. 2006, Fox and Raichle 2007). The connectivity estimates computed using brain network time courses obtained using blind decomposition techniques, such as ICA, are referred to as functional network connectivity (FNC) (Jafri et al. 2008). To capture dynamic changes of brain connectivity, one of the most common methods of estimating time-varying connectivity states is the sliding windows method in which correlation between network time courses is computed and then clustered using algorithms like k-means to identify recurring stable patterns of FNC across time and subjects (Allen et al. 2014).

Recently, there is growing interest in developing techniques to classify subjects into diagnostic groups using functional connectivity in various research domains (Allen et al. 2014, Kim et al. 2016, Plitt, Barnes, and Martin 2015, Rahaman et al. 2019, Saha et al. 2017, Saha et al. 2019). Some recent studies have evaluated classification among bipolar and schizophrenia patients using the features generated from functional connectivity (Arbabshirani et al. 2013, Shen et al. 2010, Su et al. 2013). (Shen et al. 2010) utilized an atlas-based approach to compute the mean time-courses of 116 brain regions for both resting-state controls and patients subjects. They extracted the features from each subject by computing the correlation between the time courses. After that the feature’s dimension was reduced by using dimensionality reduction method and finally, they classified patients from controls with good accuracy. Another classification approach, where features derived from sliding-window based dynamic FNC (dFNC) measures have been shown to provide significant improvements in classification accuracies of patients with schizophrenia (SZ) and bipolar disorder (BP) from healthy controls (HC) compared to sFNC measures (Rashid et al. 2016). In their work, the dFNC states were computed by clustering sliding-windowed FNC estimates using k-means with an optimum number of states obtained using elbow method (based on the average ratio of inter- and intra-cluster distances). (Byun et al. 2014) presented a classification approach where features were extracted from discrete and stable rsfMRI network states generated after applying k-means and finally, these features were used to train an SVM classifier to classify between major depressive disorder and healthy controls. The use of feature extraction from different states presents a challenge to compute the optimal number of states. Several data-driven methods to select the optimum number of centroids for k-means algorithm include: (1) the elbow method, (2) the gap statistic (Tibshirani, Walther, and Hastie 2001), and (3) the mean silhouette width (Rousseeuw 1987). The main disadvantage of elbow and silhouette methods to select the optimal number model order is that both rely on global clustering characteristics. Additionally, the elbow cannot always be unambiguously identified. Sometimes there is no elbow, or sometimes there exist several elbows in certain data distribution (Kodinariya and Makwana 2013). A limitation of the gap statistic is that it struggles to find optimum clusters when data are not separated well (Wang et al. 2018). It is unclear if the number of clusters obtained using this method is optimal for classification. To mitigate this problem, we introduce an approach that estimates the optimum number of dFNC states based on the best classification rates in a nested cross-validation analysis. While we build our current work on the previously published work by (Rashid et al. 2016), we addressed few limitations that were present in the previous work. (Rashid et al. 2016) focused on classification performance using static, dynamic, and combined connectivity features, however, the authors didn’t utilize a compact feature for classification purposes. In this current study, we explore both capturing dynamic information from time-varying connectivity windows as well as identifying a set of robust and compact features that would lead to better classification performance. Identification of the best number of clusters in k-means clustering approach that would represent the complete data matrix in the most informative way while eliminating any information redundancy is a challenging issue. This is specifically important in the case of classification algorithms where feature selection and reduction are vital for optimum model performance. Our current framework focuses on this important issue in order to capture the optimum feature sets.

In our work, we extensively consider static, dynamic, and combined (static and dynamic) features to analyze our classification task and show how the selection of optimal model order influences the classification accuracy. Meanwhile, we conduct statistical significance tested on the classification accuracies between different approaches using paired t-tests. We also investigate if further sub-clustering of the obtained k-means states (hierarchical clustering) can further improve the classification accuracy by capturing additional variation in the dFNC measures. Additionally, instead of taking all sFNC features for classification, we search for the optimum number of sFNC features that further enhances the classification accuracy when combined with dFNC features. In our final analysis, we identify the most predictive brain regions by evaluating the predictive power of sFNC features in the combined (sFNC+dFNC) classification analysis. To the best of our knowledge, our proposed method is the first approach to estimate the optimum number of dFNC states based on classification rates.

## 2. Methods

### 2.1. Participants

In our experiments, the data was acquired from the multi-site Functional Imaging Biomedical Informatics Research Network (fBIRN) project (Potkin and Ford 2009). This study included resting-state fMRI data which was scanned during the eyes-closed state and collected from 7 different sites across the USA. This data consists of 163 healthy controls (117 males, 46 females; mean age 36.9) and 151 SZ patients (114 males, 37 females; mean age 37.8), and all patients are matched by age and gender. Before scanning, informed consent was taken from each participant according to the governing internal review boards of the corresponding institutions. Previous studies from our group have used this data as well (Damaraju et al. 2014, Abrol et al. 2017).

### 2.2. Imaging parameters

Imaging data were acquired using two different types of scanner. Data from six sites were collected on a 3T Siemens Tim Trio System scanner and 3T General Electric Discovery MR750 scanner was used for one site. A standard gradient-echo echo planar imaging (EPI) paradigm consisting field of view (FOV)220 × 220 mm (64 × 64 matrices), repeat time (TR) = 2 s, echo time (TE) = 30 ms, flip angle (FA) = 77°, 162 volumes, 32 sequential ascending axial slices of 4 mm thickness and 1 mm skip was used to obtain resting-state fMRI scans.

### 2.3. Data preprocessing

Raw fMRI data went through a preprocessing pipeline using a combination of AFNI^1^, SPM^2^, and GIFT^3^ toolboxes. Briefly, the preprocessing steps included (i) subject head motion correction using the INRIAlign^4^ toolbox in SPM, (ii) slice-timing correction to address the timing differences associated with slice acquisition for rigid body motion correction, (iii) despiked the data to reduce the impact of outliers using AFNI’s 3dDespike algorithm, (iv) normalization of the data to a Montreal Neurological Institute (MNI) template and resampling to 3 mm^3^ isotropic voxels, and (v) smoothing the data to 6 mm full width at half maximum (FWHM) using AFNI’s BlurToFWHM algorithm where the smoothing was performed by a conservative finite difference approximation to the diffusion equation since this algorithm reduces scanner specific variability in smoothness by providing “smoothness equivalence” to data across multiple sites (Friedman, Hastie, and Tibshirani 2008). Finally, before group ICA, variance normalization was performed on each voxel’s time course, since this approach provides a better decomposition of subcortical sources in addition to cortical networks (Allen et al. 2010).

### 2.4. Group independent component analysis

A group independent component analysis (GICA) framework, as implemented in the GIFT software (Calhoun et al. 2001a, Erhardt et al. 2011), was used to analyze preprocessed functional data. Spatial ICA transforms the subject data into a linear mixture of spatially independent components. Principal component analysis (PCA) was used to reduce 162-time points data into 100 directions of each subject. This subject reduced data were concatenated across time and group data PCA was applied to this concatenated data to reduce it into 100 components along with the directions of maximum group variability. The Infomax algorithm (Bell and Sejnowski 1995) was used to retrieve 100 independent components from the group-PCA reduced matrix. To confirm the stability, the ICA algorithm was repeated 20 times in ICASSO^5^ and a central solution was selected using the modes of the component cluster. Finally, the spatio-temporal regression back reconstruction approach (Calhoun et al. 2001a, Erhardt et al. 2011) was used to retrieve subject-specific spatial maps (SMs) and time courses (TCs).

### 2.5. Post-ICA processing

For each subject, one sample t-test maps of each SM for all subjects were computed and thresholded to identify the regional peak activations of clusters for that component. Further, the mean power spectra of the corresponding TCs of all components were computed. A given component was identified as an intrinsic connectivity network (ICN) if that component’s peak activation clusters were on gray matter and showed less overlap with known vascular, susceptibility, ventricular, and edge regions related to head motion. Based on this selection procedure, 47 ICNs out of the 100 independent components were retained. The cluster stability/quality (Iq) index (returned from Icasso) for each estimate-cluster provides the rank for the corresponding ICA estimate. The I_q_ index for these ICNs across 20 ICASSO runs was very high (I_q_ > 0.9) for all of the components, except an ICN that resembles the language network (I_q_ = 0.74).

After obtaining 47 ICNs their corresponding TCs for each subject were detrended, orthogonalized based on subject-specific motion parameters, and then despiked to reduce the effect of outliers on subsequent FNC measures (see Supplemental Figure 1 in (Allen et al. 2014)). Despiking of corresponding TCs was performed by first detecting spikes using AFNI’s 3dDespike algorithm and replacing spikes by values getting from third-order spline fit to neighboring clean portions of the data.

**Figure 1:**
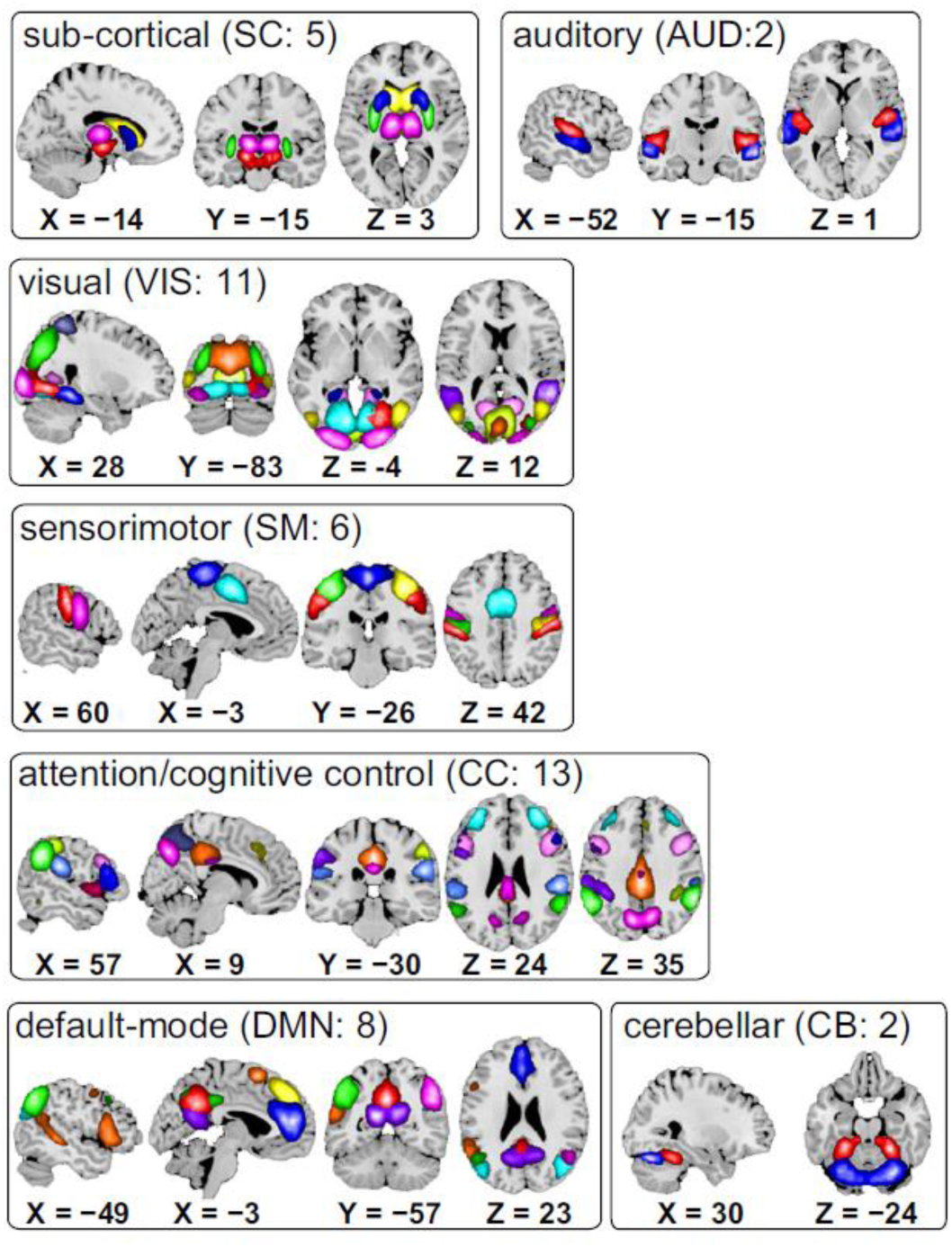
Composite maps of the 47 identified intrinsic connectivity networks (ICNs), sorted into seven subcategories. Each color in the composite maps corresponds to a different ICN

### 2.6. Static functional network connectivity (sFNC)

The pairwise correlation between ICN time courses (FNC) was computed by taking the average of connectivity among different ICNs for the duration of the scan period. The sFNC was obtained using the entire ICN time courses. Before computing FNC between ICN time courses, the ICNs were band pass filtered with a passband of [0.01–0.15] Hz using a 5^th^ order Butterworth filter. After computing the mean sFNC matrix across subjects, this matrix was arranged into modular partitions using the Louvain algorithm of the brain connectivity toolbox^6^.

Finally, the rows of sFNC matrix were partitioned into sub-cortical (SC), auditory (AUD), visual (VIS), sensorimotor (SM), a broad set of regions involved in cognitive control (CC) and attention, default-mode network (DMN) regions, and cerebellar (CB) components. To measure group differences among ICNs in sFNC, the MANCOVA framework (Allen et al. 2011) was used. Additionally, diagnosis, site, and gender were used as factors, age as a covariate, and interactions of gender and age by diagnosis. Besides, to consider residual motion related variance, mean framewise displacement (meanFD) was included as a nuisance covariate in ICA derived measures, as suggested in recent studies (Sakoglu et al. 2010, Yan et al. 2013). The reduced model is acquired by the elimination of one term and any associated interactions. At each step, the multivariate model was used to compare the variance explained in the response variable of the current full model and reduced model using the Wilks3 Lambda likelihood ratio test statistic (Christensen 2001). In this model reduction step, the term which didn’t meet the threshold value α = 0.01 for the F-test was referred to as least significant and removed.

### 2.7. Dynamic functional network connectivity (dFNC)

To compute dynamic FNC (dFNC) between two ICA time courses, a sliding window approach of window size of 22 TR (44 s) in steps of 1 TR was used by following our earlier work (Allen et al. 2014). To avoid the noise of estimated covariance using shorter-length time series, covariance was estimated from regularized inverse covariance matrix (ICOV) (Smith et al. 2011, Varoquaux et al. 2010) by graphical LASSO framework (Friedman, Hastie, and Tibshirani 2008). Besides, L1 norm constraint was applied to the inverse covariance matrix for sparsity enforcement. The regularization parameter for each subject was optimized in a cross-validation framework by assessing the log-likelihood of unseen data of the subject. The eigenvalues of estimated dynamic covariance matrices were tested as positive for validating the claim that the original graphical LASSO implementation may not certain the positive semi-definiteness of the estimated covariance matrix (Mazumder and Hastie 2012). Finally, the covariance values of the dFNC estimates for each subject were Fisher-Z transformed and residualized with respect to age, gender, and site using the reduced model which is determined from our sFNC analysis.

### 2.8. Classification framework

We evaluated our classification of static FNC, dynamic FNC and a combination of both static and dynamic FNC features following one of our earlier works (Rashid et al. 2016). In all experimental settings, we used a linear support vector machine (SVM) to compute the classification in a repeated, nested cross-validation analysis. We applied SVM on 5-fold cross-validated data with 10 repetitions. In SVM, we used a linear kernel and optimized the cost parameter (C) using the grid search approach. To find the optimal value of C, we computed cross-validated accuracy of each training fold for the different values of C [minimum C value: 0.03125, maximum C value: 2, increment size: 0.01] and used the model with optimal (i.e. validated) value of this hyperparameter to apply on the test set of that given fold.

#### 2.8.1 Static FNC approach

To perform classification on static FNC, we first split the data into five folds where each fold consists of training and testing sets. For any given fold, we split the data into training and testing datasets. We then trained a linear SVM classifier using the training data sets. Next, we validated this by classifying subjects using the testing data sets of that given fold using the trained classifier and recorded the computed accuracy. This same approach was applied to the rest of the four folds and finally, this whole procedure was repeated ten times. The conceptual framework of the sFNC approach is shown in Figure 2.

**Figure 2:**
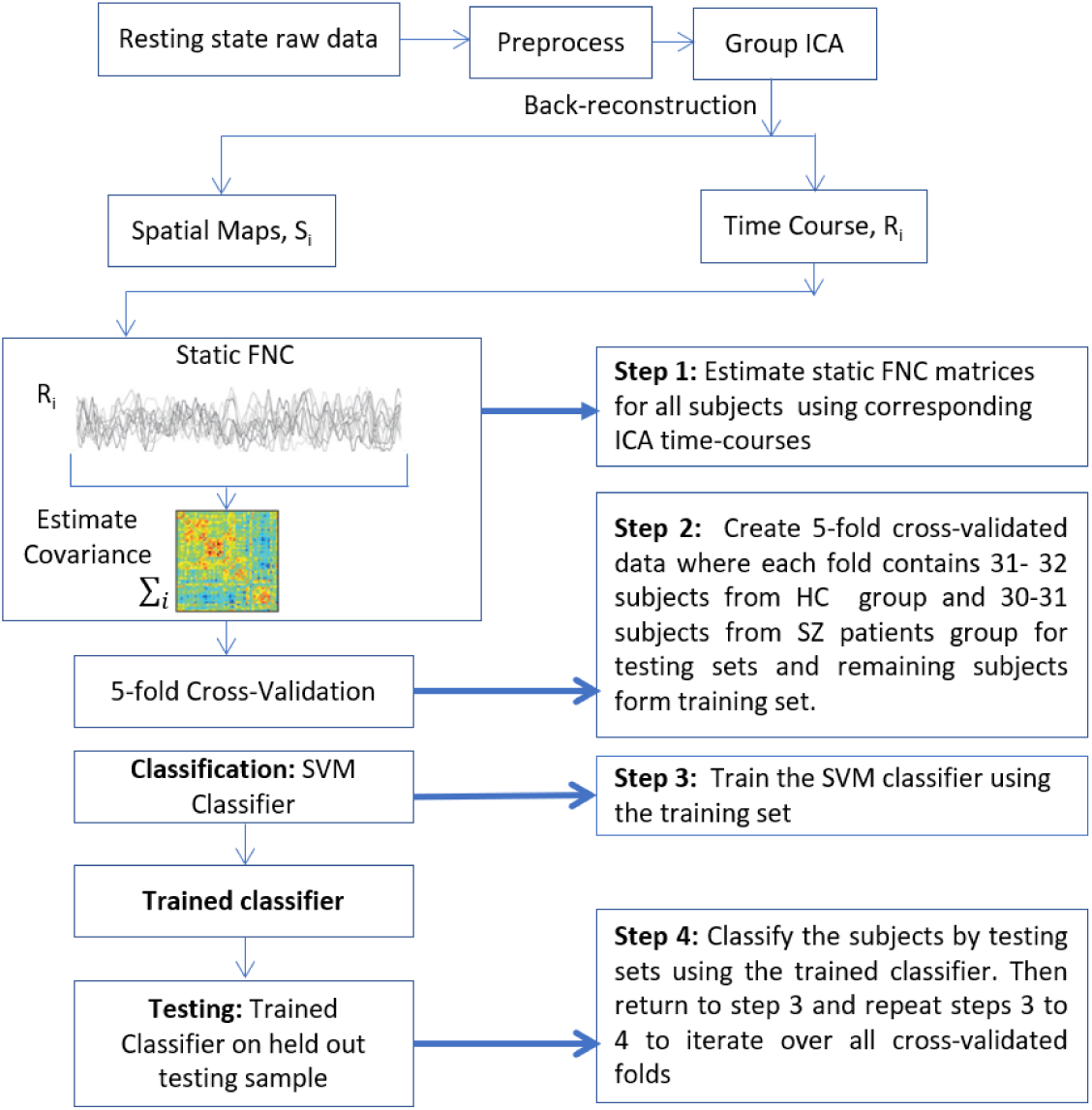
An overview of the static FNC approach. Group ICA is used to decompose resting-state data from 314 subjects into 100 independent components. Among them, 47 intrinsic connectivity networks (ICNs) are identified based on peak activation and overlapping criteria. Spatio-temporal regression back reconstruction approach is used to retrieve subject-specific spatial maps (SMs) and time courses (TCs). Static functional network connectivity is computed as the covariance of TCs which is finally used as static features.

#### 2.8.2. Dynamic FNC approach

To evaluate the classification of dynamic FNC, we used the same folds of data that are used in the static FNC approach for maintaining consistency. We applied the k-means clustering algorithm to estimate the connectivity states and further higher-level information was extracted from these states to use these as classification features. We used different model orders (model order k = 2 to 10) for k-means to extract the centroid’s information for any specific group. In other words, for any model order k and any certain repetition, we took a fold of data and applied k-means on HC and SZ groups separately. After completion of k-means, we collected centroid information from HC and SZ groups and finally concatenated these group-specific centroids to form a regression matrix **R**_groups × centroids._ Using this regression matrix, we regressed each windowed observation and computed the beta coefficient **β** (total number of beta coefficients will be, model order(k)*2 for each window). Then we computed the mean beta coefficients across all windows of any given subject which is our final dynamic FNC features Feat_**dFNC**_. After that, a linear SVM was trained by the beta coefficients of training sets and computed the accuracy for the testing sets using this trained classifier. This same approach was applied on the rest of four folds and the whole procedure was repeated ten times for any given model order. The conceptual framework of the dynamic FNC approach is shown in Figure 3.

**Figure 3:**
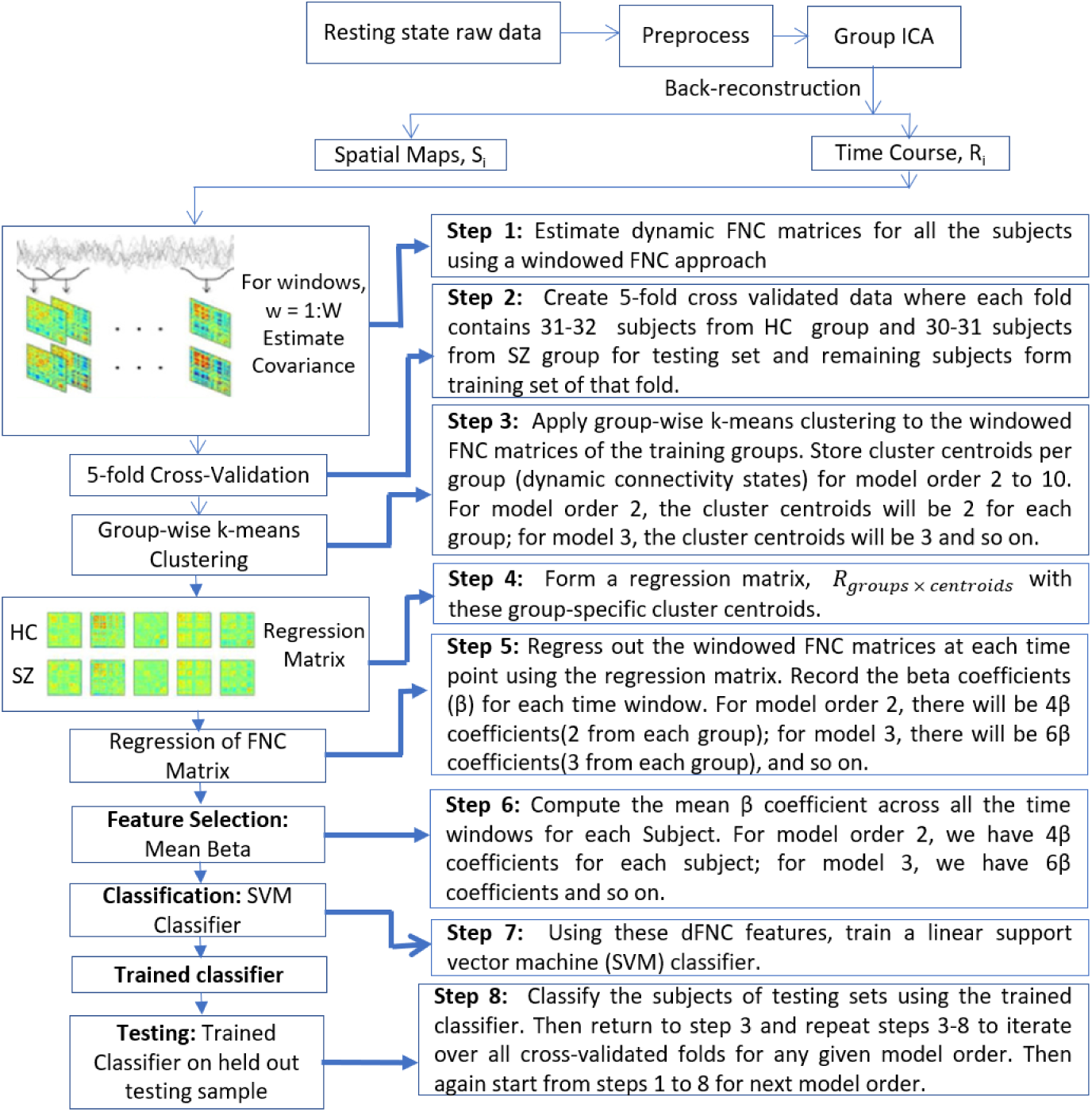
An overview of the dynamic FNC approach. At first, a group-wise k-means clustering was applied to training groups separately to obtain cluster centroids per group (HC and SZ). Next, a regression matrix was formed and the windowed FNC matrices at each time point were regressed based on this to extract the beta coefficient (**β**). Finally, the mean beta coefficient was computed across all the time windows for each subject which are considered as dynamic features

#### 2.8.3 Combined static and dynamic FNC approach

Following the same manner of static and dynamic FNC evaluations, to compute classification of the combined (static + dynamic) approach, we used the same folds of data that were used in the previous two computations. In this approach, for dimensionality reduction and feature selection, we estimated the most informative features using the double input symmetric relevance (DISR) method (Meyer and Bontempi 2006) and used these selectively reduced features instead of using the entire set of static FNC features. The DISR method selects the subset of variables (i.e. features) such that the combination of selected variables results in higher mutual information than the collective information given by each of the variables while taken individually. We ran this simulation for a range of DISR features (15, 20, 30, 40, 50, 100) to observe the fluctuation and variability of classification performance for different combinations of features. For any fold under a given model order (k), we extracted the DISR features from the sFNC training set, referring to these features as Feat_**sFNC**_. To obtain features from dynamic FNC(Feat_**dFNC**_) of the same fold of data, we applied the same approach as that used in dynamic FNC case.

In the next analysis step, the extracted static (Feat_**sFNC**_) and dynamic features (Feat_**dFNC**_) of the same subject were concatenated to form a set of combined features (Feat_sFNC+dFNC_). After that, we trained a linear SVM classifier using these combined features of the training set. For the testing set of the same fold, we used the same static features (Feat_**sFNC**_) columns that were extracted from the training sets using the DISR method. To obtain dynamic FNC features (Feat_**dFNC**_), we computed the beta coefficients **β** of each subject using the regression matrix (**R**_groups × centroids_) of the training sets following the same regression technique as outlined before. Next, we combined the static features (Feat_**sFNC**_) and dynamic features (Feat_**dFNC**_) to form a set of combined features (Feat_sFNC+dFNC_) of that testing set and classified subjects using the training set’s classifier. This same approach was applied on the rest of four folds and finally, the whole procedure was repeated 10 times for each model order. Eventually, the whole procedure was repeated for the range of the number of DISR features mentioned before. The conceptual framework of the combined (static + dynamic) FNC approach is shown in Figure 4.

**Figure 4:**
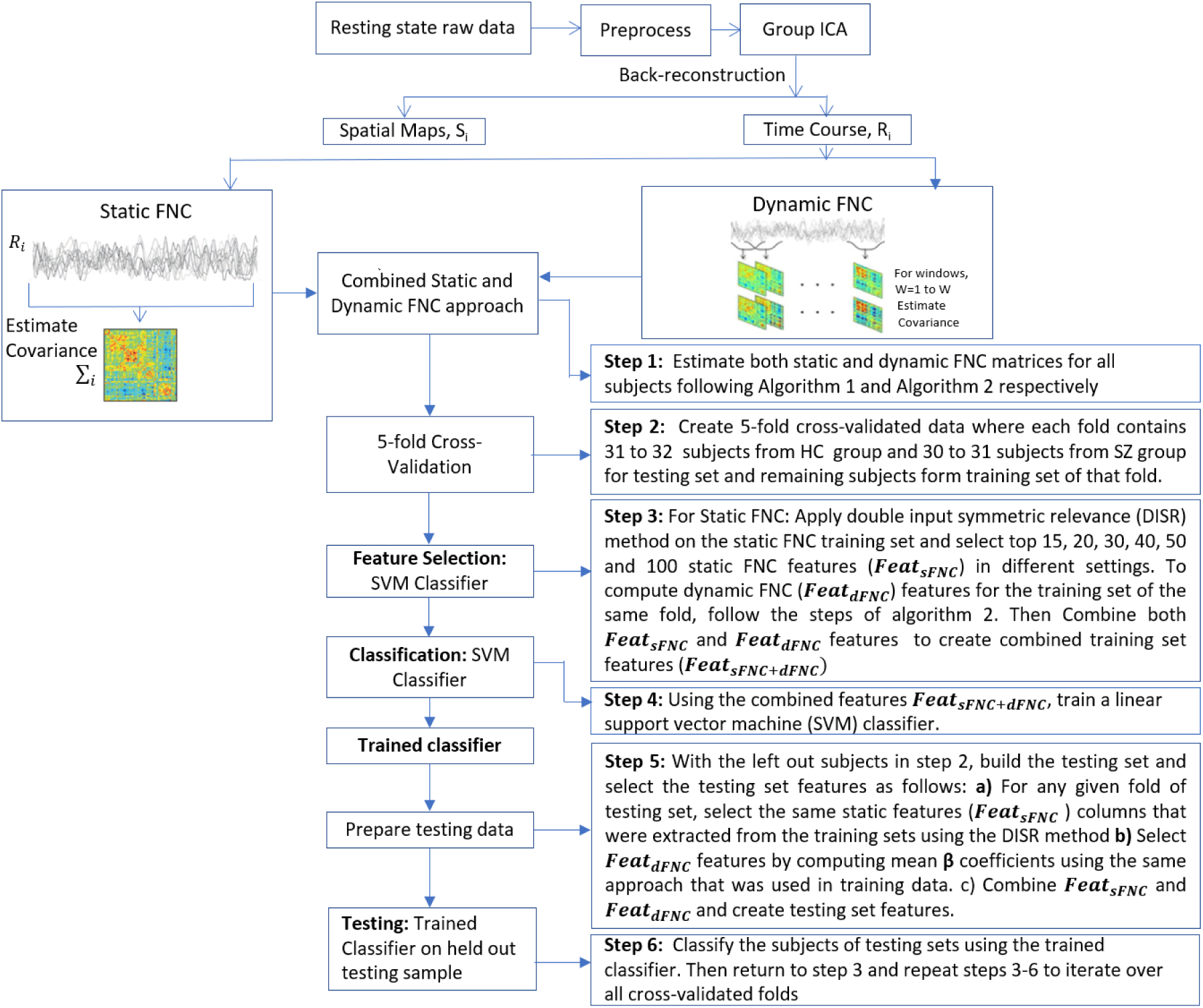
An overview of the combined (static + dynamic) approach. Here, static and dynamic features are concatenated to create the combined features for the classification. But instead of using all static features, the most informative features are selected using the double input symmetric relevance (DISR) method from pre-computed static features.

##### 2.8.3.1 Intersection approach

In this analysis, we keep all experimental setups the same as combined static and dynamic FNC analysis (section 2.8.3) except adding another layer to pick static features after applying the DISR method on static FNC data. For a given data partition, we extract 40 DISR static features for each fold and finally retain the most stable static features (Feat_**sFNC_Intersection**_) by computing the intersection across all five folds. Finally, we follow the same procedure mentioned in section 2.8.2 where we concatenate Feat_**sFNC_Intersection**_ and Feat_**dFNC**_a features and formed combined features (Feat_sFNC_Instersection+dFNC_) which is used for the classification using a linear SVM.

Additionally, to identify the dominant static features contributing to the classification accuracy throughout our whole experiments, we took the intersection of static features (Feat_**sFNC_Intersection**_) across all ten partitions of data.

#### 2.8.4 Hierarchical approach

In this experiment, we examine if further (i.e. a second level) clustering on first level clustering states can capture additional variation in dFNC measures to improve classification accuracy. Here we take information from second-level states and fit into our classification framework to investigate the contribution of this additional information by monitoring the classifier’s performance. We test this hypothesis using three different setups of experiments on the states obtained from running k-means (using the validated model order 4 only) on dynamic FNC. The reason for picking model order 4 into account for this analysis is that we get the highest accuracy for model order 4 in all of our previous experimental settings.

In setup 1, at first, we ran k-means with model order 4 on the dynamic FNC data of each fold’s training samples to estimate states. After that, k-means with model order 4 was applied again on the highest occupancy state (the state that belongs to maximum observations). Next, centroids from the first and second level states were concatenated, and beta coefficients were computed using the regression technique described in section 2.8.2.

In setup 2, at first, we ran k-means with model order 4 on the dynamic FNC data of each fold’s training samples to estimate states. After that, model order 2 was applied to the highest occupancy state. Then centroids from the first and second level states were concatenated, and beta coefficients were computed using the regression technique mentioned in section 2.8.2.

In setup 3, at first, we ran k-means with model order 4 on the dynamic FNC data of each fold’s training samples. After that, each state was further clustered by running k-means with model order 2 (resulting in 8 sub-states per fold). Finally, similar to the previous two setups, centroids from these 8 states were concatenated and the beta coefficients were computed using the regression technique mentioned in section 2.8.2.

## 3. Results

Forty-seven intrinsic connectivity networks (ICNs) were obtained through the decomposition using spatial ICA following one of our earlier investigations of whole-brain functional connectivity (Damaraju et al. 2014). Spatial maps of 47 ICNs are shown in Figure 1. The ICNs are categorized based on anatomical and functional domains including the subcortical (SC), auditory (AUD), visual (VIS), somatomotor (SM), cognitive control (CC) processes, default-mode (DMN), and cerebellar (CB) networks.

To perform classification on static FNC, we computed the pairwise correlation between the time courses of the 47 ICNs. Each subject constitutes ^47^C_2_ = 1081 brain connections; this data forms a 314 × 1081 features matrix at the group level. Note that initially we take all features of static FNC into account for classification, but for the combined (static + dynamic) case we extracted the top informative features using the DISR method as mentioned in section 2.8.3. For dynamic FNC, we computed the pairwise correlation between the time courses of 47 ICNs using the sliding-window approach (Allen et al. 2014, Rashid et al. 2014) of each dynamic window which finally formed a 314 × 136 × 1081 (subjects × windows × connections) dynamic FNC matrix. Recall that we regressed this dynamic FNC matrix by the regression matrix estimated from the group level states and that finally produce dynamic FNC features for classification as described in section 2.8.2.

The boxplots in Figures 5, 6, 7, 8, and 10 show the distribution of mean accuracies for each of the cross-validation repeats. The comparison of cross-validated classification accuracy between static and dynamic FNC is presented in Figure 5. In our classification framework, static FNC reported an overall mean accuracy of 77.16% with the range of accuracies [75.49, 79.34]. For dynamic FNC, we report a maximum accuracy for model order 4 where the mean value of the accuracy is 79.45% with range [77.96, 80.89]. The comparison metric of static vs combined (static + dynamic) for a range of the different number of DISR features is demonstrated in Figure 6. After combining static with dynamic features, the maximum accuracy was reported by model order 4 with the top 40 DISR static features where the mean accuracy is 80.45% with range [78.37, 81.87]. A similar result was observed for each case, with the same model order (k=4) being validated.

**Figure 5:**
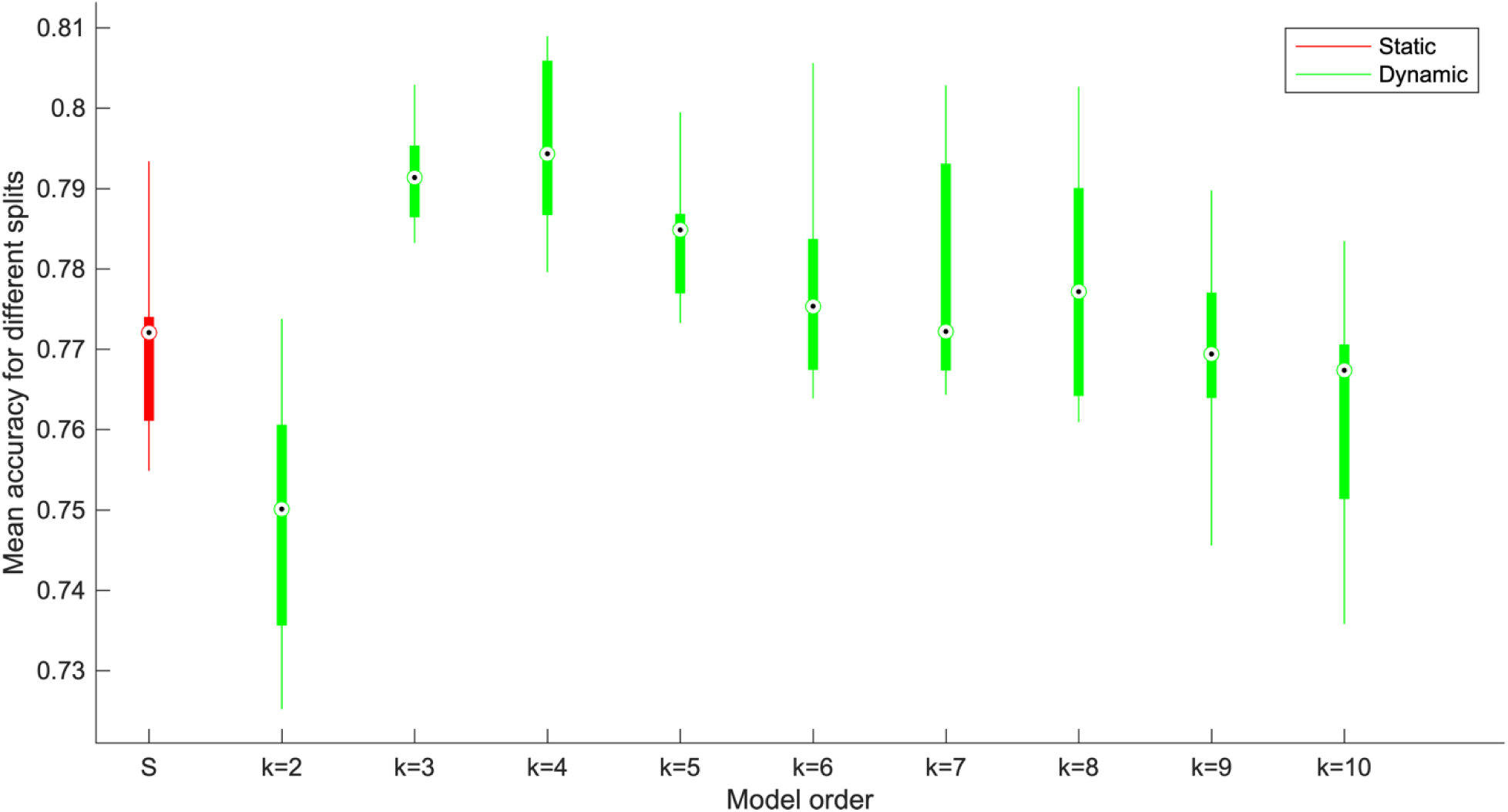
Classification accuracy of static and dynamic FNC (flat approach); On the X axis, labels k = 2-10 indicate the model order for dFNC and Y axis indicates the mean accuracy. Each boxplot consists of 10 points where each point is the computed mean accuracy of 5 folds under any given repetition of experiment (total 10 repetitions).

**Figure 6:**
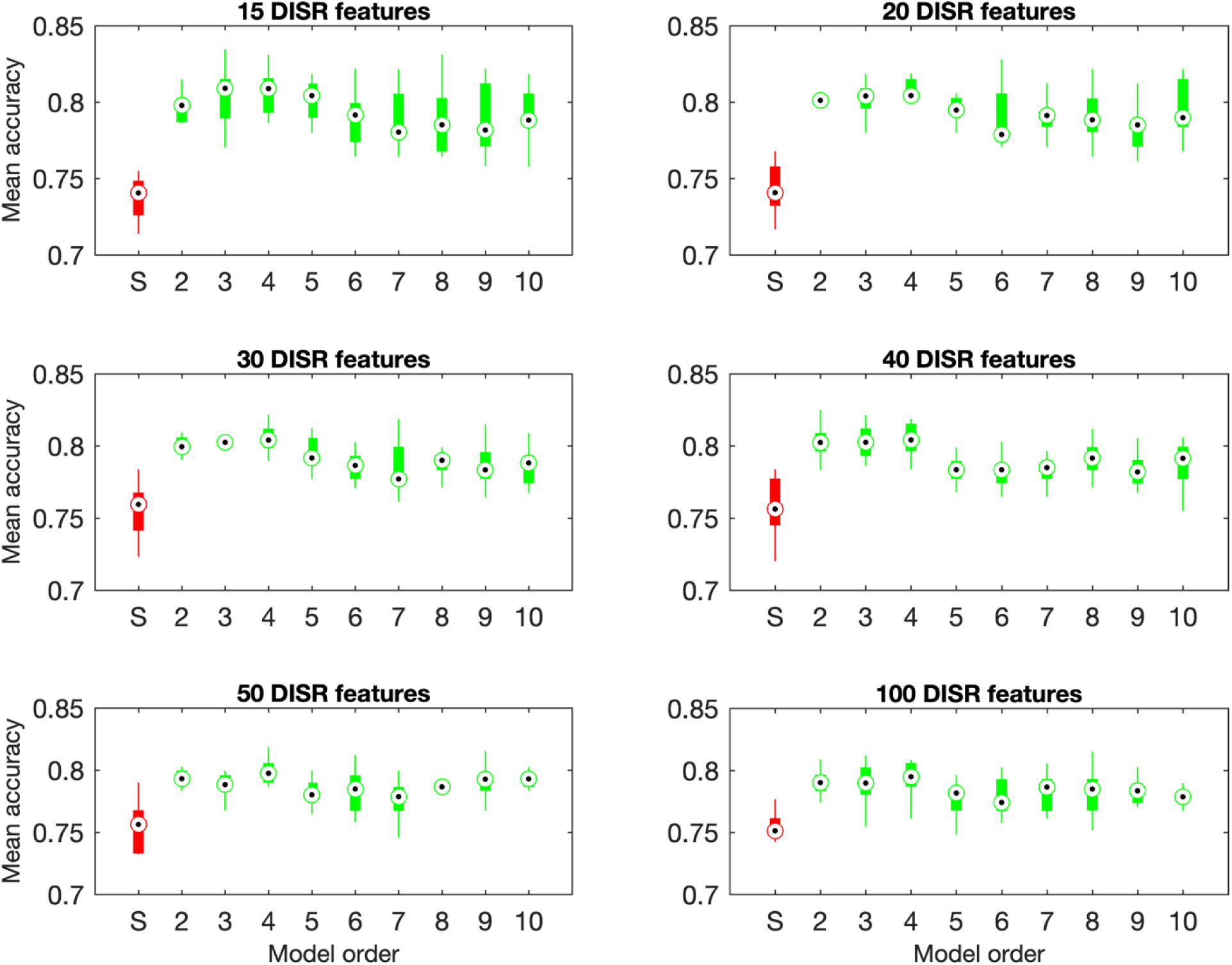
Classification accuracy of static vs combined (static + dynamic) features. Each subplot represents accuracy for different DISR features (extracted from static FNC). On the X axis, S represents accuracy for static FNC (selected static DISR features) and 2-10 indicates the model order for combined features (static DISR features + dFNC beta coefficients) and Y axis indicates the mean accuracy.

**Figure 7:**
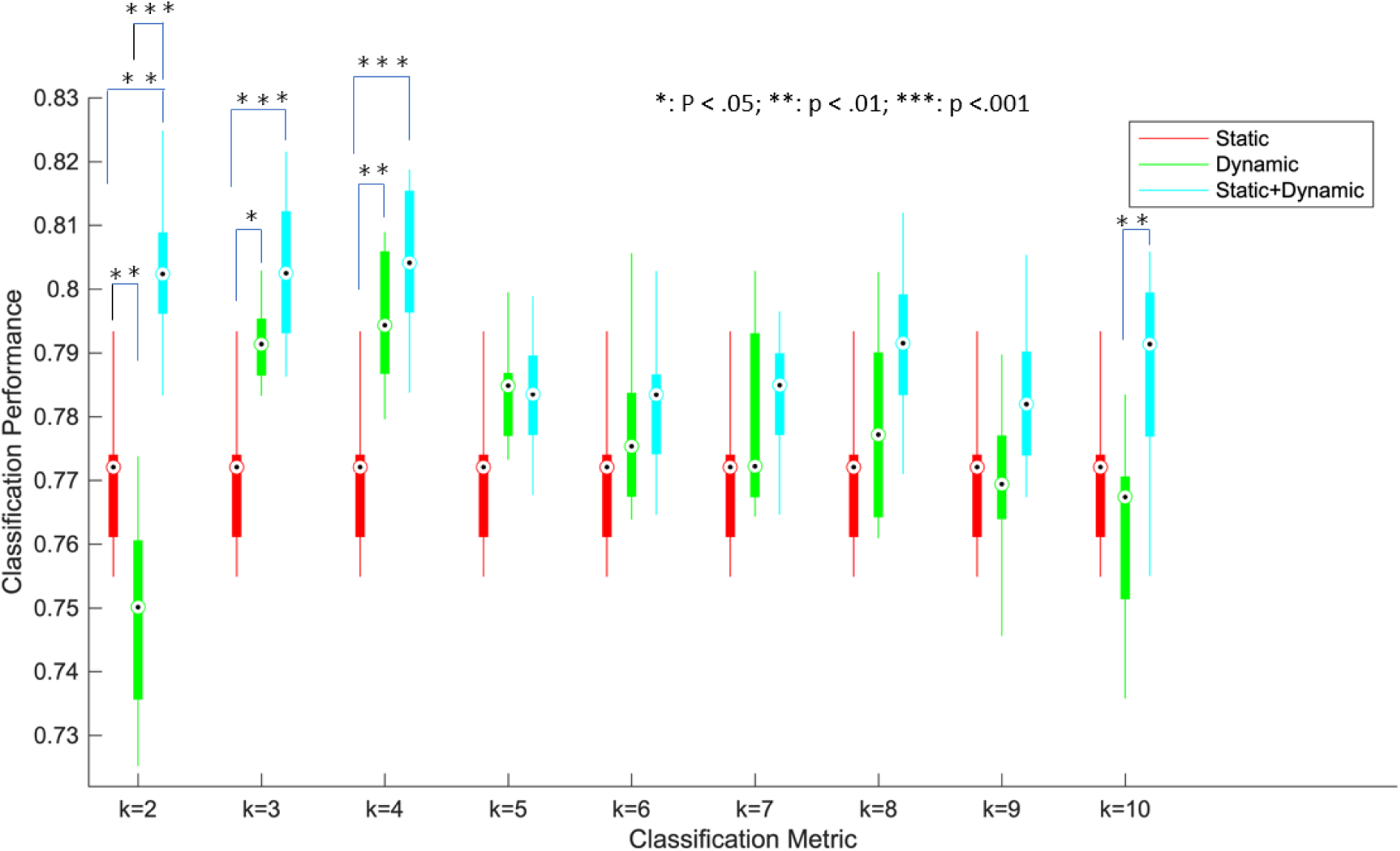
Classification accuracy of static, dynamic, and combined FNC approach. On the X axis, labels k = 2-10 indicate the model order for dFNC, and Y axis indicates the mean classification accuracy. Asterisks indicate p < 0.05 (FDR corrected), double asterisks indicate p < 0.01 (FDR corrected), Triple asterisks indicate p < 0.001 (FDR corrected).

**Figure 8:**
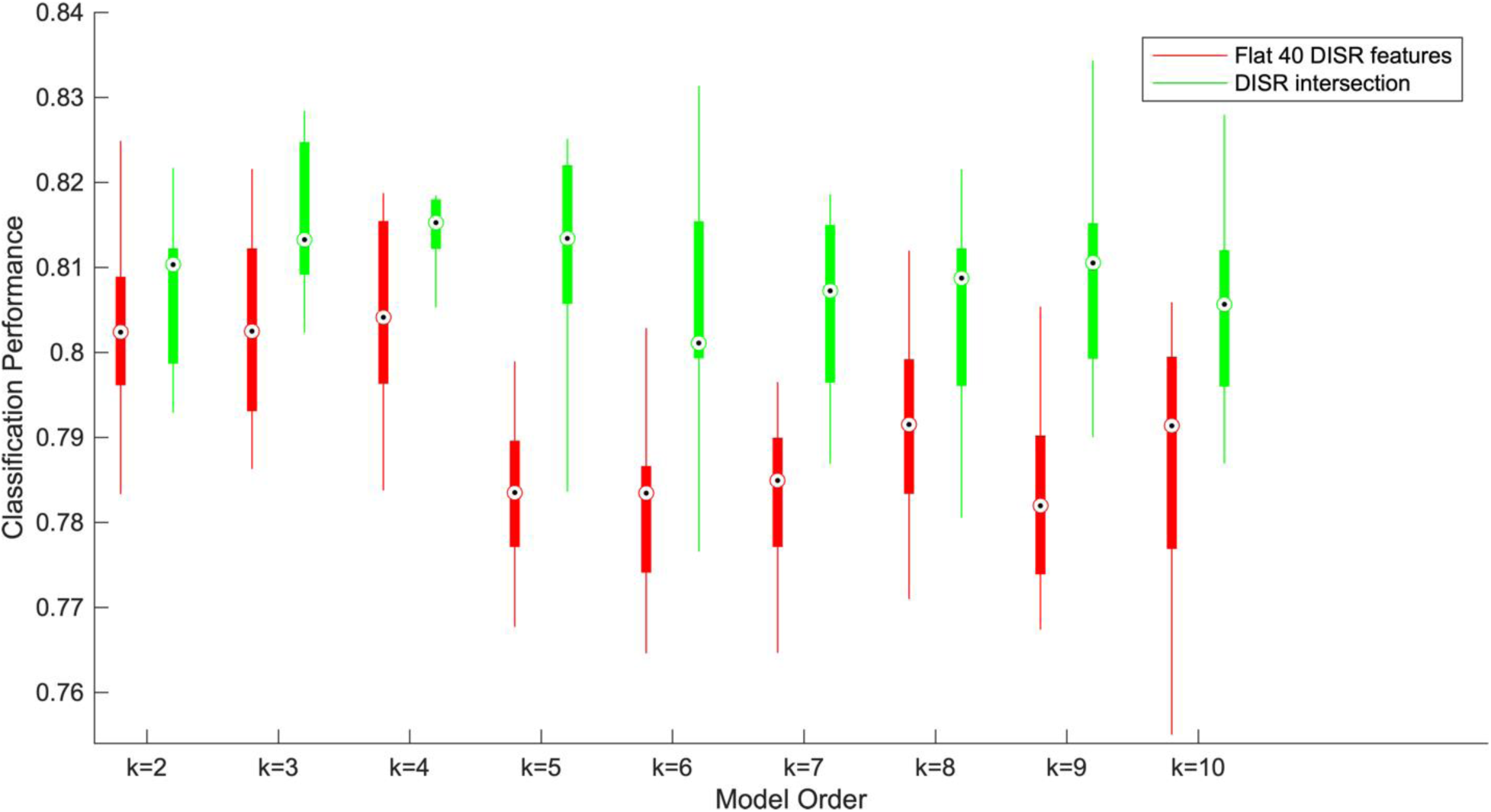
Classification accuracy of 40 DISR features vs common DISR features (obtained by intersection across all folds) of intersection approach. On X axis, labels k=2-10 indicate the model order and Y axis reports the mean accuracy values. For any model order, there exists one pair boxplot where red boxplot indicates the classification accuracy of flat 40 DISR features and the green one represents the accuracy of DISR intersection features.

We also conducted statistical significance tests on the classification accuracies for the static vs dynamic, dynamic vs combined and static vs combined approaches using paired t-tests. The results of this analysis are summarized in Figure 7. In the combined approach, we used 40 DISR static features for all model orders. Static FNC showed statistically significant improvement over dynamic FNC for model order 2. In dynamic FNC, the classification accuracy of model orders 3 and 4 showed statistically significant improvement with static FNC features. Finally, the combined (static + dynamic) approach significantly outperforms static FNC for model orders 2, 3, and 4 and dynamic FNC for model order 2 and 10. For the validated model order (k=4), we found significant improvement for the dynamic vs. static (*p* < 0.01) and combined vs. static (*p* < 0.001) cases.

The classification accuracies of the intersection approach are shown in Figure 8. In this case, the accuracy of intersected static features together with dynamic features performed slightly better than the traditional combined approach for all experimental setups. Each boxplot consists of 10 points where each point is the mean accuracy over the 5 cross-validated folds under a given partition of data. Here, yet again, the highest classification accuracy was reported for model order 4 (consistent with all previous results) where the mean accuracy is 81.53% with range [80.53, 0.8315]. Finally, we identified seven dominants static FNC features that are picked by the DISR feature selection method across all folds and all partitions as shown in Figure 9.

**Figure 9:**
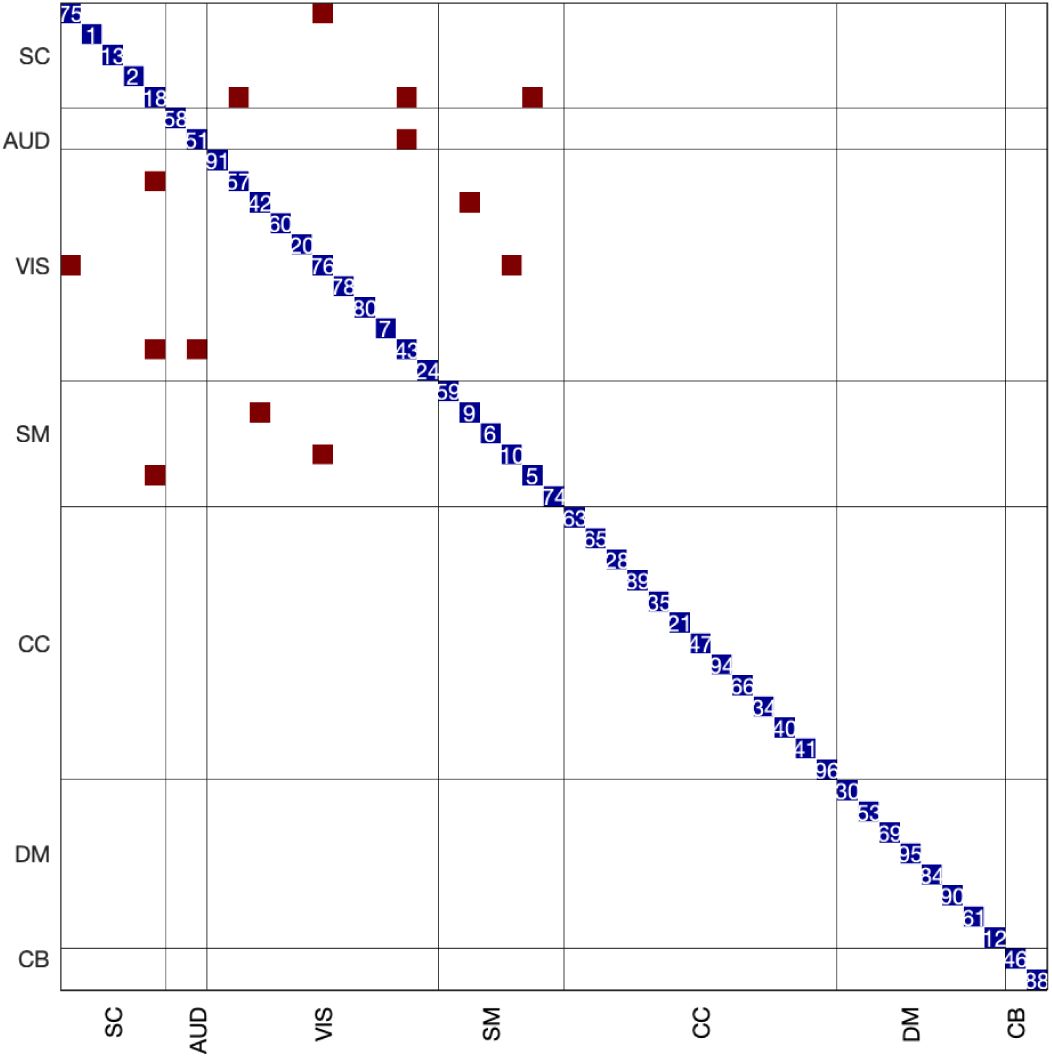
Brain mapping of most predictive static features. Three pairs of thalamocortical connections (two thalamus-visual and one thalamus-motor pair), two sensorimotor (SM)-visual (VIS) pairs, one sensorimotor (SM) - sub-cortical (SC) pair and one visual (VIS)- auditory (AUD) pair were consistently selected by the DISR method across all partitions of data. All 47 ICNs(selected from 100 independent components) are shown on the diagonal.

**Figure 10:**
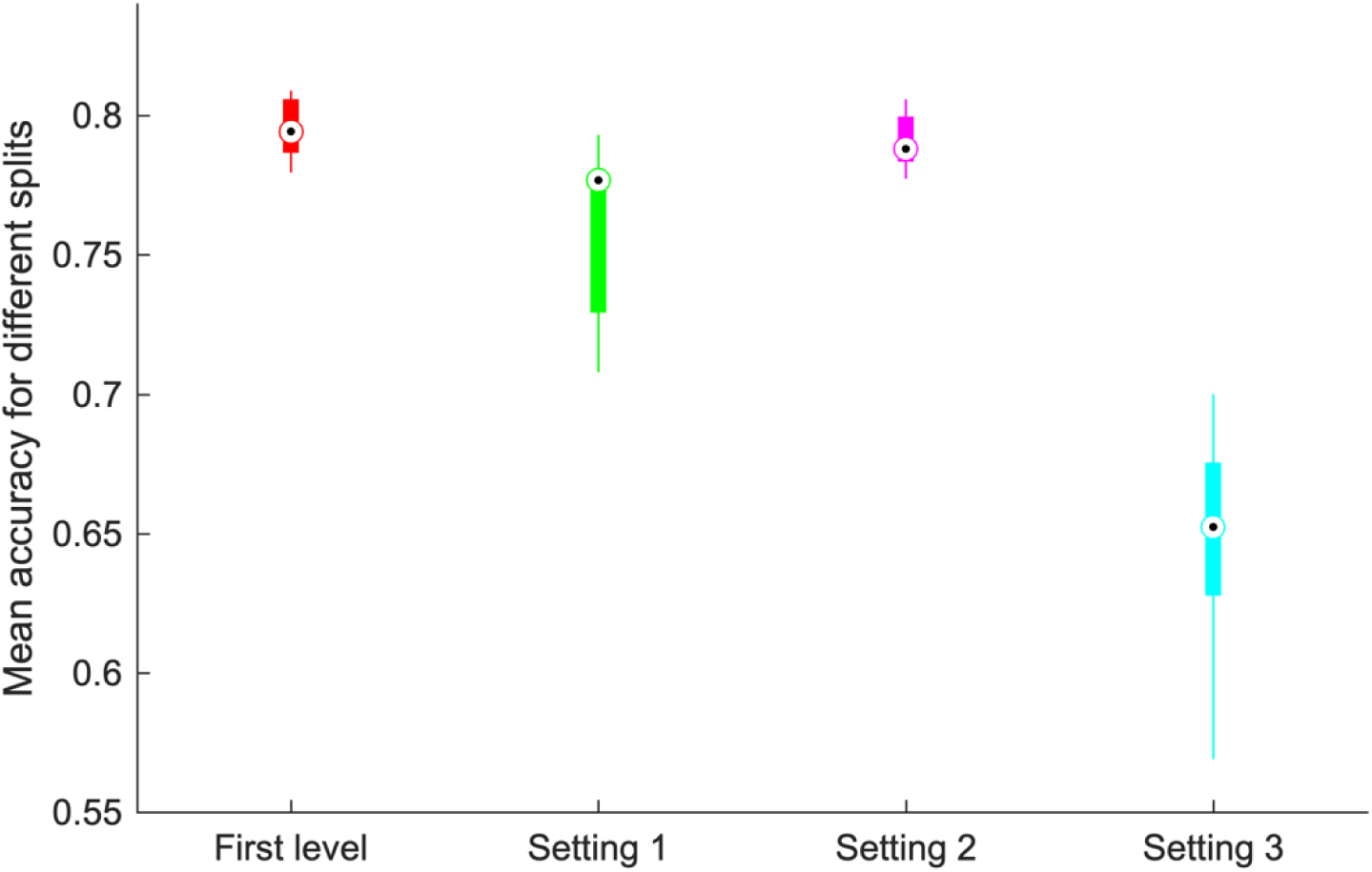
Classification accuracy of model order 4 for different settings of hierarchical approach. On X axis, “First level” indicates accuracy for model order 4 without hierarchical clustering. Settings “1-3” indicate hierarchical classification accuracy of model order 4 for three different setups.

The classification accuracies of the hierarchical approach are shown in Figure 10. In this figure, the first plot (labeled as “first level”) represents the accuracy of dynamic FNC for model order four. The other three plots highlight the classification accuracies for three different settings of second-level clustering as described in section 2.8.4. From our analysis, the classification accuracy of our traditional clustering approach always outperforms the second-level clustering settings. From model order 4 of dynamic FNC, we get maximum accuracy where the mean accuracy is 79.45% with range [77.96, 80.89]. Meanwhile, in the hierarchical approach, after using second level clustering information with model order 4 of dynamic FNC, we get maximum accuracy for Setting 2 where the mean accuracy is 79.01% with range [77.73 80.59].

## 4. Discussion

Dynamic FNC gives information about time-varying connectivity fluctuations over time (Allen et al. 2014). It captures local connectivity of each window instead of providing mean connectivity, unlike static FNC. Hence, the information that is sometimes missing in static FNC could potentially be captured by dynamic FNC approaches. A recent study has shown that these informative dynamic features help to distinguish HC and patients in a classification framework (Rashid et al. 2016). Results presented in Figure 5 show that classification accuracy of dynamic FNC outperforms static FNC for model orders 3 to 8 which is consistent with our earlier work and the best accuracy was obtained from model order 4. However, for higher model orders (k = 9, 10), the observed dynamic FNC classification accuracies drop. We speculate that this observation may due to the splitting of low occupancy states, analogous to the results observed in setting 3 in the hierarchical approach analyses (Figure 10). Also, under such circumstances, outlier clusters (for example, single-subject observations clustered together) may occur thus adding noise to the analysis. Moreover, the results demonstrate a consistent and sizable (k = 3 to 8) range of the pattern of interest in this comparison.

The classification accuracy was improved after combining the dynamic and top static FNC features together as also reported in Rashid et al. (2016). In our work, we ran experiments for different subsets of static features using the DISR method to improve the accuracy which is presented in Figure 6. Consistent with the earlier work (Rashid et al. 2016), classification accuracy improved than we obtained from static or dynamic FNC individually on separate datasets. The best accuracy was achieved for 40 DISR static FNC features in model order 4 and the combined features approach significantly outperforms static FNC for model orders 2, 3, and 4. Instead of the low accuracy of dynamic FNC compared to static FNC in model order 2, the combined approach significantly outperforms both static and dynamic FNC. A possible explanation is that the subset of static features (40 DISR features) is playing a vital role to increase the classification accuracy. This observation demonstrates that both dynamic and static FNC methods capture complementary aspects of connectivity and the classification accuracy increases after combining the dynamic and top static FNC features compared to using individual features separately.

Our analysis for identifying the most dominant static features highlighted seven brain network pairs that are consistently dominant across all 10 partitions of data as presented in Figure 9. These include three pairs of thalamocortical connections (two thalamus-visual and one thalamus-motor pair), two sensorimotor (SM) -visual (VIS) pairs, one sensorimotor (SM) - sub-cortical (SC) pair and one visual (VIS) - auditory (AUD) pair. Notably, the dysconnectivity within the thalamocortical connections has earlier been reported in (Anticevic et al. 2014) and our findings also are in line with this. The reduced anticorrelation between the thalamus and sensorimotor network in patients with SZ is a good predictor and is consistent with earlier findings (Woodward and Heckers 2016).

We further investigated hierarchical clustering states to check if information from second level sub-states can contribute to improving accuracy. Our results (Figure 10) show that classification accuracy after including the second-level information didn’t outperform the first-level clustering results. Based on this classification accuracy, it can be concluded that there is not enough variability in dFNC measures at the second-level of clustering to improve classification accuracy. We speculate that this observation may be for the splitting of low occupancy states and the information from these hierarchical states may not be significant to improve classification accuracy.

From all our experiments, we achieve the best classification accuracy for model order 4 (i.e. k = 4 in k-means clustering) to distinguish HC and SZ subjects of the fBIRN data. However, the traditional elbow method suggested the optimal value of k = 5 for this data (Damaraju et al. 2014). This suggests that the optimal model order of k-means algorithm for classification can be different from model order obtained from other standard methods. Thus, our work presents a strong case to conduct an exhaustive search for validating the clustering model order in a similar classification, prediction, or characterization brain imaging applications.

## 5. Limitations and future directions

We consider, as dynamic features, the mean regression coefficients across time for each of the derived HC or SZ states, when fitted on the sliding window-based dynamic functional connectivity samples. While the mean regression coefficients provide a compact and meaningful feature sets, there are other possible approaches to extract dynamic connectivity features that should be considered in future work. For instance, instead of considering the mean strategy for extracting dynamic features, other approaches like the standard deviation or the instantaneous change in similarity to a given state might be interesting work to explore in the future. There are some experimental limitations to choose the optimal window size in the dynamic FNC approach. The optimal window size should be able to measure the variability of functional connectivity and capture any short time effects (Sakoglu et al. 2010). We used a sliding window approach of fixed window size of 22 TR (44 s) in steps of 2 TR following one of our earlier works (Allen et al. 2014, Damaraju et al. 2014). Capturing different patterns of variability using different window lengths in a dynamic FNC approach and finally fitting the features into the classification framework can be an interesting work for further research.

In addition to the linear SVM classifier, a variety of other classifiers can be further used to examine the robustness of our proposed approach. In SVM we used a linear kernel and optimized the parameter “C” using a grid search approach. To the best of our knowledge, this is the first study to propose a classification approach for model order selection for clustering in a dynamic FNC analysis. A detailed comparison of different kernels of SVM and optimization of different SVM hyperparameters would be interesting to explore in the future.

## Acknowledgment

This study was supported by NIH R01EB020407, R01MH118695.

## Footnotes

1 http://afni.nimh.nih.gov/.

2 http://www.fil.ion.ucl.ac.uk/spm/.

3 http://mialab.mrn.org/software/gift/index.html.

4 http://www-sop.inria.fr/epidaure/Collaborations/IRMf/INRIAlign.html

5 http://research.ics.aalto.fi/ica/icasso/

6 https://sites.google.com/site/bctnet/

